# Novavax NVX-COV2373 triggers potent neutralization of Omicron sub-lineages

**DOI:** 10.1101/2022.07.14.500148

**Authors:** Jinal N. Bhiman, Simone I. Richardson, Bronwen E. Lambson, Prudence Kgagudi, Nonkululeko Mzindle, Haajira Kaldine, Carol Crowther, Glenda Gray, Linda-Gail Bekker, Novavax trial clinical lead author group, Vivek Shinde, Chijioke Bennett, Gregory M. Glenn, Shabir Madhi, Penny L. Moore

**Affiliations:** National Institute for Communicable Diseases of the National Health Laboratory Services, Johannesburg, South Africa; MRC Antibody Immunity Research Unit, School of Pathology, University of the Witwatersrand, Johannesburg, South Africa; The South African Medical Research Council, Tygerberg, South Africa; Institute of Infectious Disease and Molecular Medicine, University of Cape Town, Cape Town, South Africa; The Desmond Tutu HIV Centre, University of Cape Town, Cape Town, South Africa; Novavax, Inc, Gaithersburg, Maryland, United States; South African Medical Research Council Vaccines and Infectious Diseases Analytics Research Unit, Faculty of Health Sciences, University of the Witwatersrand, Johannesburg, South Africa; Centre for the AIDS Programme of Research in South Africa, University of Kwazulu-Natal, Durban, South Africa; Wits Vaccines and Infectious Diseases Analytics (VIDA) Research Unit; Wits RHI, Faculty of Health Sciences, University of the Witwatersrand, Johannesburg, South Africa; Limpopo Clinical Research Initiative; Soweto Clinical Trials Centre (SCTC); Centre for Lung Infection and Immunity, Division of Pulmonology, Department of Medicine and UCT Lung Institute & South African MRC/UCT Centre for the Study of Antimicrobial Resistance, University of Cape Town, Cape Town, South Africa; Faculty of Infectious and Tropical Diseases, Department of Immunology and Infection, London School of Hygiene and Tropical Medicine, London, UK; South African Tuberculosis Vaccine Initiative (SATVI), Department of Pathology, Institute of Infectious Disease and Molecular Medicine and Division of Immunology, Faculty of Health Sciences, University of Cape Town, Observatory, Cape Town; Setshaba Research Centre (SRC); Josha Research; Verulam and Isipingo Clinical Research Site, South African Medical Research Council, HIV and other Infectious Diseases Research Unit (HIDRU); Durban International Clinical Research Site, Enhancing Care Foundation; Madibeng Centre for Research; The Aurum Institute; Pretoria Clinical Research Centre; KwaPhila Health Solution; Peermed CTC (PTY) - MERC Kempton; Mzansi Ethical Research Centre Middleburg

## Abstract

The SARS-CoV-2 Omicron (B.1.1.529) Variant of Concern (VOC) and its sub-lineages (including BA.2, BA.4/5, BA.2.12.1) contain spike mutations that confer high level resistance to neutralizing antibodies. The NVX-CoV2373 vaccine, a protein nanoparticle vaccine, has value in countries with constrained cold-chain requirements. Here we report neutralizing titers following two or three doses of NVX-CoV2373. We show that after two doses, Omicron sub-lineages BA.1 and BA.4 were resistant to neutralization by 72% (21/29) and 59% (17/29) of samples. However, after a third dose of NVX-CoV2373, we observed high titers against Omicron BA.1 (GMT: 1,197) and BA.4 (GMT: 582), with responses similar in magnitude to those triggered by three doses of an mRNA vaccine. These data are of particular relevance as BA.4 is emerging to become the dominant strain in many locations, and highlight the potential utility of the NVX-CoV2373 vaccine as a booster in resource-limited environments.

## Main text

The SARS-CoV-2 Omicron (B.1.1.529) Variant of Concern (VOC)^1^ and its sub-lineages^2^ (including BA.2, BA.4, BA.5, BA.2.12.1) contain changes to the spike driven by immune escape, and are relatively immune evasive compared with ancestral-like virus to neutralizing antibodies elicited by coronavirus disease 2019 (COVID-19) vaccines^3,4^. Similarly, individuals infected with SARS-CoV-2 exhibit reduced neutralizing titers against multiple Omicron sub-lineages^3^. Neutralization escape by the Omicron VOC has also been observed following vaccination, regardless of the vaccine type and platform^3–8^, including with two doses of the NVX-CoV2373 vaccine^9^. However, booster doses, especially using mRNA vaccines, enhance neutralization capacity against Omicron^4,7^. The NVX-CoV2373 vaccine, which was tested in two phase 3 trials in the US, UK and Mexico demonstrated 90% efficacy against symptomatic and 100% efficacy against severe COVID-19^10,11^. A Phase 2b trial in South Africa in 2020-2021 demonstrated 48% efficacy against symptomatic infection COVID-19, likely due to relatively antibody-evasive neutralization resistant Beta variant, despite 100% efficacy against severe disease^12^. The vaccine has received authorization for use by the European Medicines Agency and is listed on the World Health Organization’s emergency use listing for COVID-19 vaccines^12–14^. This protein-based vaccine is appealing in low-and middle-income countries (LMICs) because of its stability and reduced cold chain requirements. Here, we investigated the effect of a third dose on the neutralizing capacity of NVX-CoV2373 vaccinee sera.

We tested neutralization of the ancestral D614G, Beta, Omicron BA.1 and Omicron BA.4/BA.5 by NVX-CoV2373 vaccinee sera following a 2 dose (n = 29) and 3 dose (n = 48) regimen. Fourteen days after two doses of NVX-CoV2373, geometric mean titers (GMT) were highest against the D614G variant (GMT: 1,401), with reductions in GMT to 173 (8.1-fold reduction), 34 (41-fold reduction) and 47 (30-fold reduction) against Beta, Omicron BA.1 and Omicron BA.4/BA.5 respectively. The Omicron sub-lineages BA.1 and BA.4/BA.5 were resistant to neutralization, with titers less than limit of detection of the assay, by 72% (21/29) and 59% (17/29) of samples after the 2^nd^ dose of vaccine (**Fig 1**, grey).

**Figure 1.**
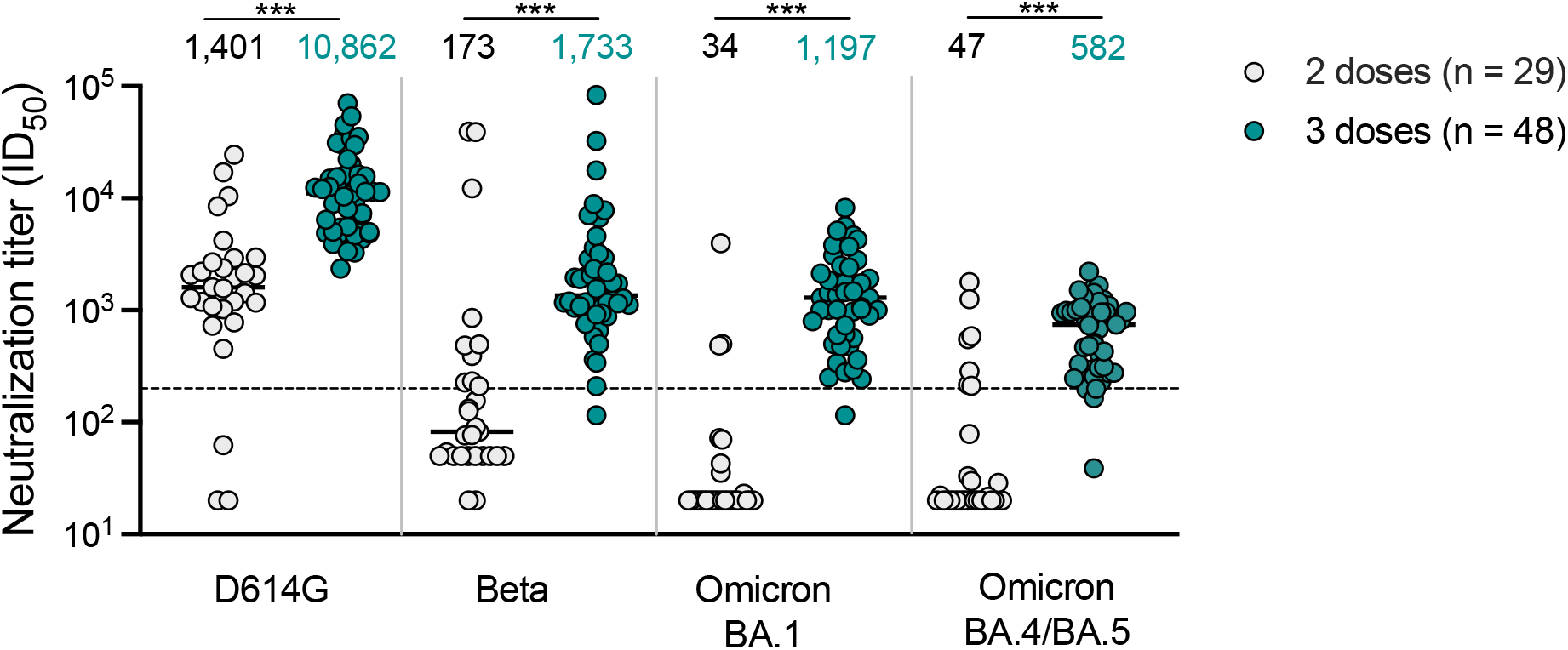
Neutralization of SARS-CoV-2 variants by NVX-CoV2373 vaccinee plasma. Neutralization of ancestral D164G, Beta, Omicron BA.1 and Omicron BA.4/BA.5 pseudoviruses by NVX-CoV2373 vaccinee plasma following 2 (grey) or 3 (teal) doses. Geometric mean titers (GMT) for each virus are shown above the individual points, and percent of specimens where no neutralization was observed (red) is indicated in the pie charts. Number of vaccinee specimens tested are indicated and p values were calculated using the Mann-Whitney t-test for non-parametric data with p < 0,001 for D614G, Beta, Omicron BA.1 and Omicron BA.4/BA.5. Dashed line indicates the neutralization level at 20.2% of the mean convalescent level (ID_50_ = 200), which provides an estimated 50% protection against detectable SARS-CoV-2 infection per the analysis by Khoury *et al^15^*. Samples were used at a starting dilution of 1 in 20 (limit of detection) with a seven 3-fold dilutions to create a titration series.

At one month after the third dose of the NVX-CoV2372 vaccine, neutralizing antibody activity with titers above 1:200 were evident against the Beta and Omicron BA.1 variants in all but one sample (n=47/48). The neutralizing antibody titers against the D614G variant were boosted to GMT of 10,862. Furthermore, we observed a significant 10-,35- and 12-fold increase in titers against Beta (GMT: 1,733), Omicron BA.1 (GMT: 1,197) and Omicron BA.4/BA.5 (GMT: 582) respectively (**Fig 1,** teal), though titers were 6- to 18-fold lower than those against D614G. Convalescent plasma titers (**Supp Fig** 1) were used to relate Omicron sub-lineage titres to 50% protection levels as described by Khoury et al^15^, and showed that after three doses of NVX-CoV2373, all but one sample exceeded this threshold.

We next compared neutralization of Omicron BA.1 and BA.4/BA.5 following multi-dose regimens of adenoviral, mRNA and protein-based vaccines. As expected, 2 doses of AD26.COV2.S elicited 10- and 14-fold lower GMT against BA.1 than 3 doses of the BNT162b2 and NVX-CoV2373 vaccines respectively (**Fig 2**). Similarly the AD26.COV2.S vaccine elicited 12- and 11-fold lower GMT against BA.4/BA.5 than 3 does of either the BNT162b2 and NVX-CoV2373 vaccines. Most third dose BNT162b2 and NVX-CoV2373 plasma were able to neutralize Omicron BA.1 and BA.4/BA.5 at titers greater than 200, while only 13-38% of the AD26.COV2.S samples achieved these titers. Given the lower GMT for all three boosted vaccine regimens against BA.4/BA.5, it was unsurprising that 91% (44/48) NVX-CoV2373 and 83% (10/12) BNT162b2 samples, compared to 13% (1/8) AD26.COV.S samples neutralized at titres above 1 in 200 threshold. The NVX-CoV2373 third dose plasma GMT against BA.1 and BA.4/BA.5 was comparable to BNT162b2, with the BA.1 data trending higher for NVX-COV2373.

**Figure 2.**
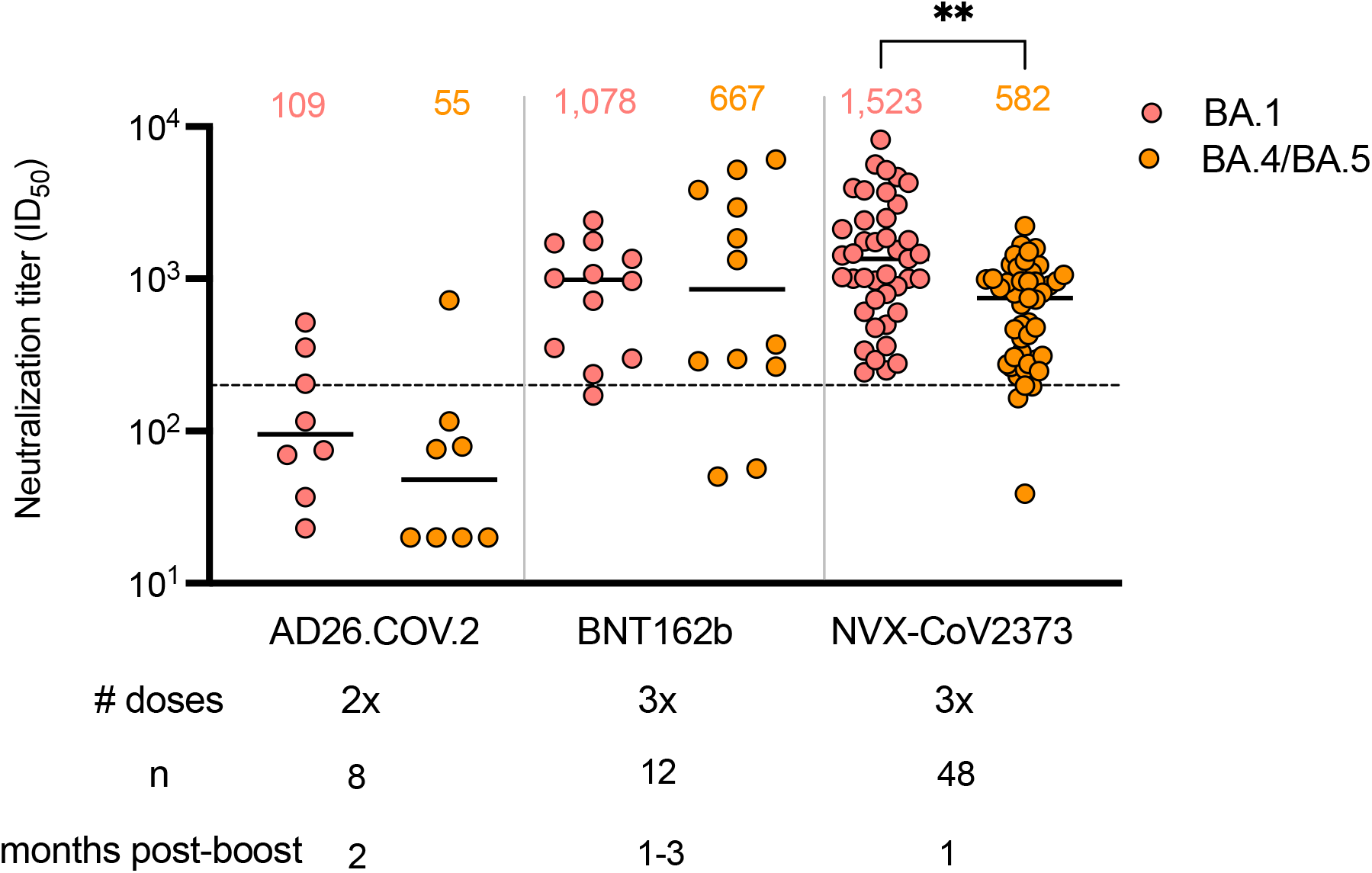
Neutralization of Omicron BA.1 and BA.4/BA.5 by boosted vaccinee plasma. Neutralization of Omicron BA.1 and BA.4/BA.5 by vaccinee plasma following 2 doses of the AD26.COV2S or 3 doses of the BNT162b2 or NVX-CoV2373 vaccines. Number of doses, number of samples and date of sample collection after boost for each group are indicated. Geometric mean titers (GMT) for each virus are shown above the individual points, P values were calculated using two-way ANOVA with p < 0,001 for AD26CoV2.S versus NXV-CoV2373 and p = 0,0011 for NVX-CoV2373 BA.1 versus BA.4/BA.5). Dashed line indicates the neutralization level at 20,2% of the mean convalescent level (ID_50_ = 200), which provides an estimated 50% protection against detectable SARS-CoV-2 infection per the analysis by Khoury *et* al^15^.

In summary, we report enhanced neutralization of Omicron BA.1 and BA.4/BA.5 following three doses of the NVX-CoV2373 vaccine with responses comparing well to an mRNA vaccine. We note that six months after two doses of NVX-CoV2373, increased binding antibodies were reported, and responses may mature further^10^. As durability of vaccine platforms varies, future studies should assess this for NVX-CoV2373 neutralization at later time-points^9^.

## Acknowledgments

We thank Dr Thandeka Moyo-Gwete, Brent Oosthuysen, Donald Mhlanga, Frances Ayres, Haajira Kaldine, Nelia P. Manamela, Sebotsana Rasebotsa, Sharon Madzorera, Thanusha Naidoo, Thopisang Motlou for technical assistance in generating plasmids and / or proteins for this study.

## Author contributions

Designed the study, performed analyses and wrote the manuscript: JNB, PLM Performed experiments and analysed data: SIR, BEL, PK NM, HK, Data curation and project management: CC PIs for Sisonke (AD26CoV2.S) Trial: GG, LGB Site PIs for Novavax Trial: AK, LFa, LFo, QB, KD, MT, MM,ZH, NS, SH, MA, CL, CG, UL, NJ, GK PIs for Novavax Trial: VS, CB, GMG SM

## Methods

### Samples and ethics approvals

Individuals vaccinated with two or three doses of the NVX-CoV2373 vaccine were sampled at 14 days after the second dose or 35 days after the third dose. This trial is registered under the ClinicalTrials.gov number, NCT04533399, and the protocol was approved by the South African Health Products Regulatory Authority and by the institutional review board at each trial centre as described in detail by Shinde and colleagues^12^. Health care workers vaccinated with two dose of AD26.COV2.S (5 × 10^10^ viral particles) as part of the Sisonke implementation trial were sampled at 2 months after vaccination. This trial is registered under the ClinicalTrials.gov number, NCT05148845, and the protocol was approved by the South African Health Products Regulatory Authority. These Sisonke individuals were recruited at the National Institute for Communicable Diseases (NICD), Johannesburg. Individuals vaccinated with two and three doses of the BNT162b22 vaccine were sampled at 2 months after the second dose or 1-3 months after the third dose and were recruited from Johannesburg. This study was given ethics approval by the University of the Witwatersrand Human Research Ethics Committee (Medical) M210465. All individuals provided written informed consent.

### Lentiviral pseudovirus production and neutralization assay

The 293T/ACE2. MF cells modified to overexpress human ACE2 were kindly provided by M. Farzan (Scripps Research). Cells were cultured in DMEM (Gibco BRL Life Technologies) containing 10% heat-inactivated fetal bovine serum (FBS) and 3μgml-1 puromycin at 37 °C, 5% CO2. Cell monolayers were disrupted at confluency by treatment with 0.25% trypsin in 1mM EDTA (Gibco BRL Life Technologies). The SARS-CoV-2, Wuhan-1 spike, cloned into pCDNA3.1 was mutated using the QuikChange Lightning Site-Directed Mutagenesis kit (Agilent Technologies) to include D614G (ancestral D164G) or L18F,D80A, D215G, Δ242-244, K417N, E484K, N501Y, D614G, A701V (Beta) or Δ69-70, T915I, Δ143-145, Δ211, L212I, ins 214 EPE, G339D, S371L, S373P, S375F, K417N, N440K, G446S, S477N, T478K, Q493R, G496S, Q498R, N501Y, Y505H, T547K, D614G, N679K, P681H, N764K, D796Y, N856K, Q954H, N969K, L981F (Omicron BA.1) or T19I, L24S, Δ25-27, Δ69-70, G142D, V213G, G339D, S371F, S373P, S375F, T376A, D405N, R408S, K417N, N440K, L452R, S477N, T478K, E484A, F486V, Q498R, N501Y, Y505H, D614G, H655Y, N679K, P681H, N764K, D796Y, Q954H, N969K (Omicron BA.4/BA.5). Pseudoviruses were produced by co-transfection with a lentiviral backbone (HIV-1 pNL4.luc encoding the firefly luciferase gene) and either of the SARS-CoV-2 spike plasmids with PEIMAX (Polysciences). Culture supernatants were clarified of cells by a 0.45-μM filter and stored at −80 °C. Plasma samples were heat-inactivated and clarified by centrifugation. Pseudovirus and serially diluted plasma/sera were incubated for 1h at 37 °C, 5% CO2. Cells were added at 1×10^4^ cells per well after 72h of incubation at 37 °C, 5% CO2, luminescence was measured using PerkinElmer Life Sciences Model Victor X luminometer. Neutralization was measured as described by a reduction in luciferase gene expression after single-round infection of 293T/ACE2.MF cells with spike-pseudotyped viruses. Titers were calculated as the reciprocal plasma dilution (ID50) or monoclonal antibody concentration (IC50) causing 50% reduction of relative light units. Equivalency was established through participation in the SARS-CoV-2 Neutralizing Assay Concordance Survey Concordance Survey 1 run by EQAPOL and VQU, Duke Human Vaccine Institute. Cell-based neutralization assays using live virus or pseudovirus have demonstrated high concordance, with highly correlated 50% neutralization titers (Pearson r=0.81–0.89). Per Khoury and colleagues^15^, we used a threshold neutralization titer of 1 in 200, which is at 20.2% (**Fig** 1, dashed line) that of the convalescent plasma mean level (GMT: 993), estimated to provide a 50% level of protection from infection.

**Supplementary Figure 1.**
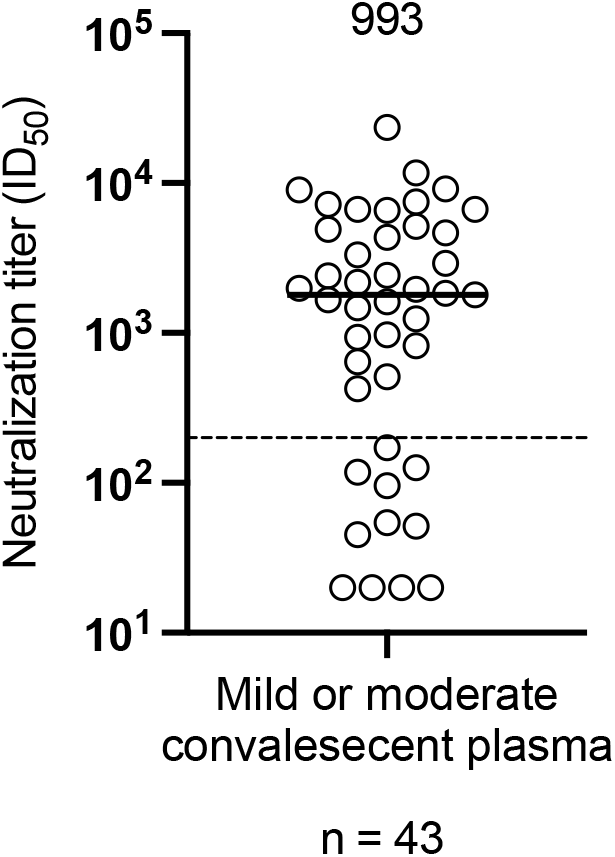
Neutralization of SARS-CoV-2 D614G by convalescent plasma. Neutralization of ancestral D164G pseudoviruses by convalescent plasma collected during the study period of the NVX-CoV2373 trial in South Africa. Geometric mean titer (GMT) is shown above the individual points, and number of specimens tested are indicated. Dashed line indicates the neutralization level at 20,2% of the mean convalescent level, which provides an estimated 50% protection against detectable SARS-CoV-2 infection per the analysis by Khoury *et* al^15^.

**Supplementary Table 1.** Vaccinee metadata

| **Study ID** | **Age** | **Gender** | **Vaccine received** | **Days post last vaccine dose** | **Months post last vaccine dose** |
| --- | --- | --- | --- | --- | --- |
| 939 | 50-59 | F | AD26.COV.S | - | 2 |
| 940 | 30-39 | F | AD26.COV.S | - | 2 |
| 945 | 30-39 | F | AD26.COV.S | - | 2 |
| 953 | 30-39 | F | AD26.COV.S | - | 2 |
| 952 | 20-29 | F | AD26.COV.S | - | 2 |
| 947 | 40-49 | F | AD26.COV.S | - | 2 |
| 950 | 30-39 | F | AD26.COV.S | - | 2 |
| 955 | 30-39 | F | AD26.COV.S | - | 2 |
| 956 | 20-29 | F | AD26.COV.S | - | 2 |
| 948 | 50-59 | F | AD26.COV.S | - | 2 |
| 930 | 60-69 | F | BNT162b2 | - | 2 |
| 931 | 60-69 | F | BNT162b2 | - | 2 |
| 936 | 70-79 | F | BNT162b2 | - | 2 |
| 933 | 70-79 | F | BNT162b2 | - | 2 |
| 928 | 70-79 | M | BNT162b2 | - | 2 |
| 929 | 70-79 | F | BNT162b2 | - | 2 |
| 932 | 60-69 | M | BNT162b2 | - | 2 |
| Pf_3_2 | 50-59 | F | BNT162b2 | - | 1 |
| Pf_3_3 | 30-39 | M | BNT162b2 | - | 1 |
| Pf_3_4 | 40-49 | F | BNT162b2 | - | 3 |
| Pf_3_1 | 40-49 | F | BNT162b2 | - | 3 |
| ZA0010017 | 20-29 | M | NVX-CoV2373 | 35 | - |
| ZA0010041 | 30-39 | M | NVX-CoV2373 | 35 | - |
| ZA0010042 | 30-39 | M | NVX-CoV2373 | 35 | - |
| ZA0010093 | 30-39 | M | NVX-CoV2373 | 14 or 35 | - |
| ZA0010106 | 20-29 | F | NVX-CoV2373 | 14 or 35 | - |
| ZA0010126 | 30-39 | F | NVX-CoV2373 | 35 | - |
| ZA0010170 | 40-49 | M | NVX-CoV2373 | 35 | - |
| ZA0010258 | 30-39 | M | NVX-CoV2373 | 35 | - |
| ZA0010263 | 30-39 | F | NVX-CoV2373 | 35 | - |
| ZA0010325 | 30-39 | F | NVX-CoV2373 | 14 or 35 | - |
| ZA0010331 | 20-29 | M | NVX-CoV2373 | 35 | - |
| ZA0010332 | 30-39 | M | NVX-CoV2373 | 35 | - |
| ZA0010337 | 20-29 | F | NVX-CoV2373 | 14 or 35 | - |
| ZA0010340 | 20-29 | M | NVX-CoV2373 | 14 or 35 | - |
| ZA0010351 | 50-59 | F | NVX-CoV2373 | 14 or 35 | - |
| ZA0010358 | 30-39 | F | NVX-CoV2373 | 14 or 35 | - |
| ZA0010424 | 30-39 | F | NVX-CoV2373 | 14 or 35 | - |
| ZA0010440 | 20-29 | F | NVX-CoV2373 | 35 | - |
| ZA0010452 | 30-39 | M | NVX-CoV2373 | 35 | - |
| ZA0010469 | 30-39 | M | NVX-CoV2373 | 14 or 35 | - |
| ZA0010516 | 30-39 | F | NVX-CoV2373 | 35 | - |
| ZA0010549 | 30-39 | M | NVX-CoV2373 | 35 | - |
| ZA0010557 | 20-29 | F | NVX-CoV2373 | 35 | - |
| ZA0010577 | 18-19 | F | NVX-CoV2373 | 35 | - |
| ZA0010611 | 40-49 | F | NVX-CoV2373 | 35 | - |
| ZA0010615 | 20-29 | F | NVX-CoV2373 | 35 | - |
| ZA0010636 | 20-29 | F | NVX-CoV2373 | 35 | - |
| ZA0010644 | 30-39 | F | NVX-CoV2373 | 35 | - |
| ZA0010656 | 20-29 | M | NVX-CoV2373 | 35 | - |
| ZA0030061 | 20-29 | M | NVX-CoV2374 | 14 or 35 | - |
| ZA0030077 | 30-39 | M | NVX-CoV2375 | 14 or 35 | - |
| ZA0030117 | 40-49 | M | NVX-CoV2376 | 14 or 35 | - |
| ZA0030118 | 30-39 | M | NVX-CoV2377 | 14 or 35 | - |
| ZA0030138 | 30-39 | M | NVX-CoV2378 | 14 or 35 | - |
| ZA0030187 | 20-29 | M | NVX-CoV2379 | 14 or 35 | - |
| ZA0030199 | 30-39 | M | NVX-CoV2380 | 14 or 35 | - |
| ZA0030204 | 20-29 | F | NVX-CoV2381 | 14 or 35 | - |
| ZA0120014 | 20-29 | M | NVX-CoV2382 | 14 or 35 | - |
| ZA0120048 | 30-39 | M | NVX-CoV2383 | 14 or 35 | - |
| ZA0150166 | 18-19 | M | NVX-CoV2384 | 14 or 35 | - |
| ZA0150234 | 18-19 | M | NVX-CoV2385 | 14 or 35 | - |
| ZA0150252 | 20-29 | M | NVX-CoV2386 | 14 or 35 | - |
| ZA0150327 | 20-29 | M | NVX-CoV2387 | 14 or 35 | - |
| ZA0180009 | 50-59 | F | NVX-CoV2388 | 14 or 35 | - |
| ZA0180024 | 20-29 | M | NVX-CoV2389 | 35 | - |
| ZA0180058 | 20-29 | M | NVX-CoV2390 | 14 or 35 | - |
| ZA0180080 | 20-29 | F | NVX-CoV2391 | 35 | - |
| ZA0180081 | 30-39 | M | NVX-CoV2392 | 14 or 35 | - |
| ZA0180128 | 18-19 | M | NVX-CoV2393 | 35 | - |
| ZA0180138 | 20-29 | M | NVX-CoV2394 | 14 or 35 | - |
| ZA0180224 | 18-19 | M | NVX-CoV2395 | 14 or 35 | - |
| ZA0180293 | 20-29 | M | NVX-CoV2396 | 35 | - |
| ZA0180306 | 20-29 | F | NVX-CoV2397 | 35 | - |
| ZA0180371 | 20-29 | M | NVX-CoV2398 | 35 | - |
| ZA0180424 | 18-19 | M | NVX-CoV2399 | 35 | - |
| ZA0180434 | 20-29 | F | NVX-CoV2400 | 35 | - |
| ZA0180481 | 20-29 | F | NVX-CoV2401 | 35 | - |
| ZA0180513 | 20-29 | M | NVX-CoV2402 | 35 | - |
| ZA0180518 | 40-49 | F | NVX-CoV2403 | 35 | - |
| ZA0180529 | 20-29 | M | NVX-CoV2404 | 35 | - |
| ZA0180532 | 30-39 | M | NVX-CoV2405 | 35 | - |
| ZA0180589 | 30-39 | F | NVX-CoV2406 | 35 | - |
| ZA0180611 | 20-29 | M | NVX-CoV2407 | 35 | - |
| ZA0180690 | 20-29 | F | NVX-CoV2408 | 35 | - |
| ZA0200005 | 20-29 | M | NVX-CoV2409 | 35 | - |
| ZA0200040 | 20-29 | M | NVX-CoV2410 | 35 | - |
| ZA0200044 | 20-29 | M | NVX-CoV2411 | 14 or 35 | - |
| ZA0200055 | 30-39 | F | NVX-CoV2412 | 35 | - |
| ZA0200062 | 30-39 | M | NVX-CoV2413 | 35 | - |
| ZA0200118 | 20-29 | F | NVX-CoV2414 | 35 | - |
| ZA0200125 | 20-29 | M | NVX-CoV2415 | 35 | - |
| ZA0210015 | 20-29 | F | NVX-CoV2416 | 35 | - |
| ZA0210206 | 30-39 | F | NVX-CoV2417 | 35 | - |
| ZA0210380 | 20-29 | F | NVX-CoV2418 | 35 | - |

